# Antibiotic treatment protocols revisited: The challenges of a conclusive assessment by mathematical modeling

**DOI:** 10.1101/372938

**Authors:** Hildegard Uecker, Sebastian Bonhoeffer

## Abstract

Hospital-acquired bacterial infections lead to prolonged hospital stays and increased mortality. The problem is exacerbated by antibiotic resistant strains that delay or impede effective treatment. To ensure a successful therapy and to manage antibiotic resistance, treatment protocols that draw on several different antibiotics might be used. This includes the administration of drug cocktails to individual patients (“combination therapy”) but also the random assignment of drugs to different patients (“mixing”) and a regular switch in the default drug used in the hospital from drug A to drug B and back (“cycling”). For the past 20 years, mathematical models have been used to assess the prospects of antibiotic combination therapy, mixing, and cycling. But while tendencies in their ranking across studies have emerged, the picture remains surprisingly inconclusive and incomplete. In this article, we review existing modeling studies and demonstrate by means of examples how methodological factors complicate the emergence of a consistent picture. These factors include the choice of the criterion by which the effects of the protocols are compared, the model implementation, and its analysis. We thereafter discuss how progress can be made and suggest future modeling directions.

## Introduction

For many decades, bacterial infections have been successfully treated with antibiotics, making formerly life-threatening diseases easily treatable. However, the rapid evolution of resistance and the slow discovery of new antimicrobial compounds increasingly reduce treatment options. In the European Union, resistant bacteria are responsible for about 25, 000 deaths per year as estimated by the World Health Organization (Fact sheet “Antibiotic resistance”, October 2015). While drastic restrictions in the use of antibiotics are urgently needed to stop this alarming trend, antibiotics must be used as wisely as possible whenever their application is required. Unfortunately, it is far from obvious to know what is wise, and we need to understand what the consequences of different treatment strategies are to be able to make more rational choices.

While unable to replace empirical research and clinical trials, mathematical models have helped to gain insight into the effects of antimicrobial stewardship. Mathematical studies profit from several strengths. They rely on explicit and well-defined assumptions, allow us to explore ideas much faster than clinical trials, and are not subject to practical and ethical restrictions. A question that has been repeatedly addressed in theoretical studies over the past 20 years concerns the integrated application of multiple antibiotics across a community – usually a hospital ward – during the phase of empirical therapy (Table 1). The idea is that strains that are resistant to one of the drugs are suppressed by the other. Within a single patient, this can be achieved by the administration of two (or more) antibiotics in combination (“combination therapy”). Across a community, it is also an option to prescribe different drugs to different patients in order to create a heterogeneous environment for the bacteria. Most prominently, the default drug can be cycled in time (“cycling”), creating temporal heterogeneity, or a fraction of the patients can receive each drug (“mixing”), creating spatial heterogeneity. The use of two (or more) antibiotics in either form, however, comes at the risk of selecting for double resistant strains that can withstand both drugs. Identifying which strategy best treats infections in the face of resistance and at the same time selects least for multiply resistant bacteria is challenging. In practice, the increased risk of side effects and higher economic costs associated with combination therapy are additional factors but current theoretical work only assesses the disease dynamics and emergence of resistance under the various strategies.

**Table 1:**
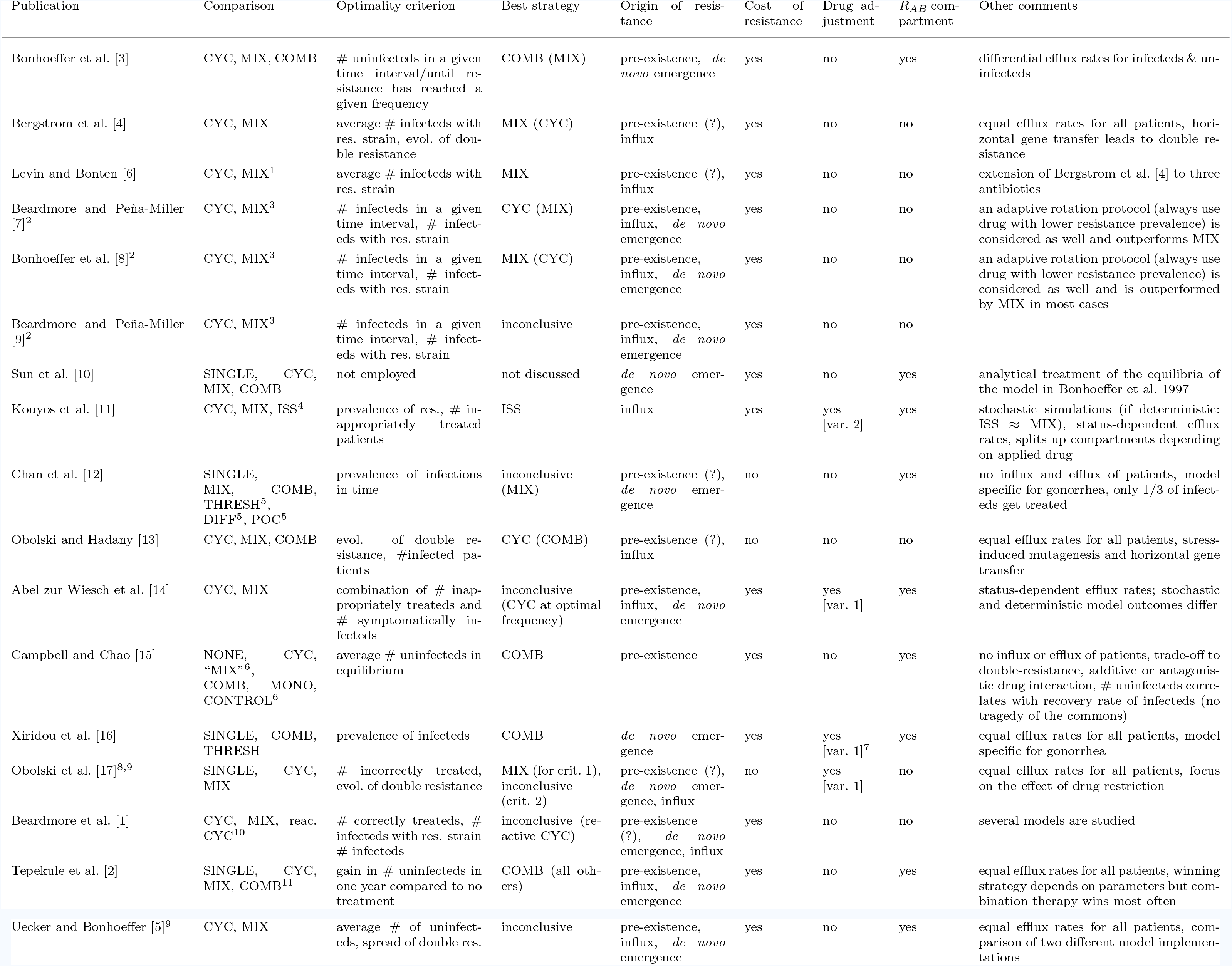
Literature overview. Best strategy: We list the strategy that emerges as the best one overall. A strategy that might win under some circumstances is added in brackets. When the picture as a whole remains inconclusive but a strategy seems to have some advantage over the others, we note this strategy in brackets. Origin of resistance: Pre-existence of resistance refers to the initial conditions at time 0. If not specified, we put a question mark. Drug adjustment: Patients are switched to a working drug either through a switch from drug A to drug B or vice versa (variant 1) or through switch to a narrow-spectrum antibiotic (variant 2). 1. Three antibiotics are used.
2. Bonhoeffer et al. [8] is a response to Beardmore and Peña-Miller [7], and Beardmore and Peña-Miller [9] is a response to Bonhoeffer et al. [8].
3. Drugs can be used unequally (i.e. different proportions in MIX; different periods of use in CYC), and for both strategies, the performance under optimal drug use are compared.
4. ISS stands for informed switching strategy. The antibiotic for incoming patients depends on the prevalence of resistance to both drugs in the hospital, and several variants of ISS are tested.
5. For THRESH, drug A is used until resistance has reached a threshold. For DIFF, different strategies are used depending on the risk group, defined through the rate of partner change. For POC, point-of-care testing is available such that resistant infections can be identified and treated accordingly.
6. Mixing is different here since each half of the population cycles the drugs. For CONTROL, evolution of resistance is impossible (we exclude it from the comparison).
7. Patients infected with a double resistant strain can be re-treated with a third antibiotic, to which resistance can evolve as well.
8. The main text of the article focuses on the effect of restricted vs an equal use of a third antibiotic. We only consider the briefly investigated two-drug model from supplementary information S3.
9. These two studies do not aim to generally assess the performance; we report the results for the examples shown in the articles.
10. Reactive cycling denotes a strategy where the default drug is always the one for which resistance is currently less prevalent.
11. Reactive versions of cycling and mixing are included as well.

Modeling studies tend to rank the three principal treatment protocols in the order “combination therapy > mixing ≥ cycling” but the picture is not conclusive (see Table 1). Since no strategy is optimal under all circumstances [1, 2], we cannot expect to find a universally true ranking. However, the picture might become more integral through a better understanding of the conditions that favor one or the other strategy. In this article, we pinpoint the difficulties that mathematical studies face in the assessment of antimicrobial treatment protocols. To do so, we set up a model for the spread of bacterial infections within a hospital that follows the traditional modeling approach. It combines features of the two original models by Bonhoeffer et al. [3] and Bergstrom et al. [4], similar to the model in Tepekule et al. [2]. We choose to explicitly refer to a hospital setting because the framework is most relevant for antibiotic treatment in hospitals but for most of the manuscript, this is only a choice of wording. We then demonstrate step by step how the ranking of strategies is affected by factors other than the biology of the pathogen. The most important one is the choice of the optimality criterion by which the performance of a treatment protocol is assessed. Due to the complexity of the problem that requires the consideration of several aspects, multiple criteria are in use, making study outcomes difficult to compare.

But also purely technical aspects can pose obstacles in arriving at congruent conclusions. Last, we discuss how these problems might be addressed and how current models could be extended, helping mathematical models to better meet their potential in assessing antimicrobial treatment protocols.

## The modeling framework

Two of the first mathematical studies on the topic were conducted by Bonhoeffer et al. [3] and Bergstrom et al. [4]. Considerable work has built on this early research, which we summarize in Table 1 (see also supplementary information section S1). While differing in many respects, almost all existing studies are based on the same general approach. They follow the number of uninfected and infected patients over time, where infected patient populations are divided up into several classes according to the infecting bacterial strain (which is characterized by the resistance profile).

The examples in the present article are based on an overarching model that incorporates the most fundamental processes, which are influx and efflux of patients, infection, clearance, the *de novo* emergence of resistance, and replacement infection, but it does not incorporate more detailed features such as explicitly modeled drug interactions [2, 5]. It brings together elements from Bonhoeffer et al. [3] and Bergstrom et al. [4], as detailed in SI section S1. As all studies except for the short commentary article by Levin and Bonten [6], we consider the use of two (and not more) drugs. All issues raised in the present article carry over to future models that would incorporate more drugs. (There is no a priori reason to assume that the ranking of strategies is independent of the number of drugs used.) The same applies to models considering other strategies such as informed cycling strategies that make use of information on the prevalence of resistance [7, 11].

A flow diagram of the model is shown in Figure 1. Patients can be uninfected or infected by one of four bacterial strains – the sensitive strain that responds to both drugs, the two strains that are resistant to only one drug, and the double resistant strain. Following the convention in the field, we denote the number of uninfected patients by *X*, the number of patients infected by the sensitive strain by *S*, and the number of patients infected by a resistant strain by *R_•_*, where the subscript indicates to which drug(s) the strain in resistant.

**Figure 1:**
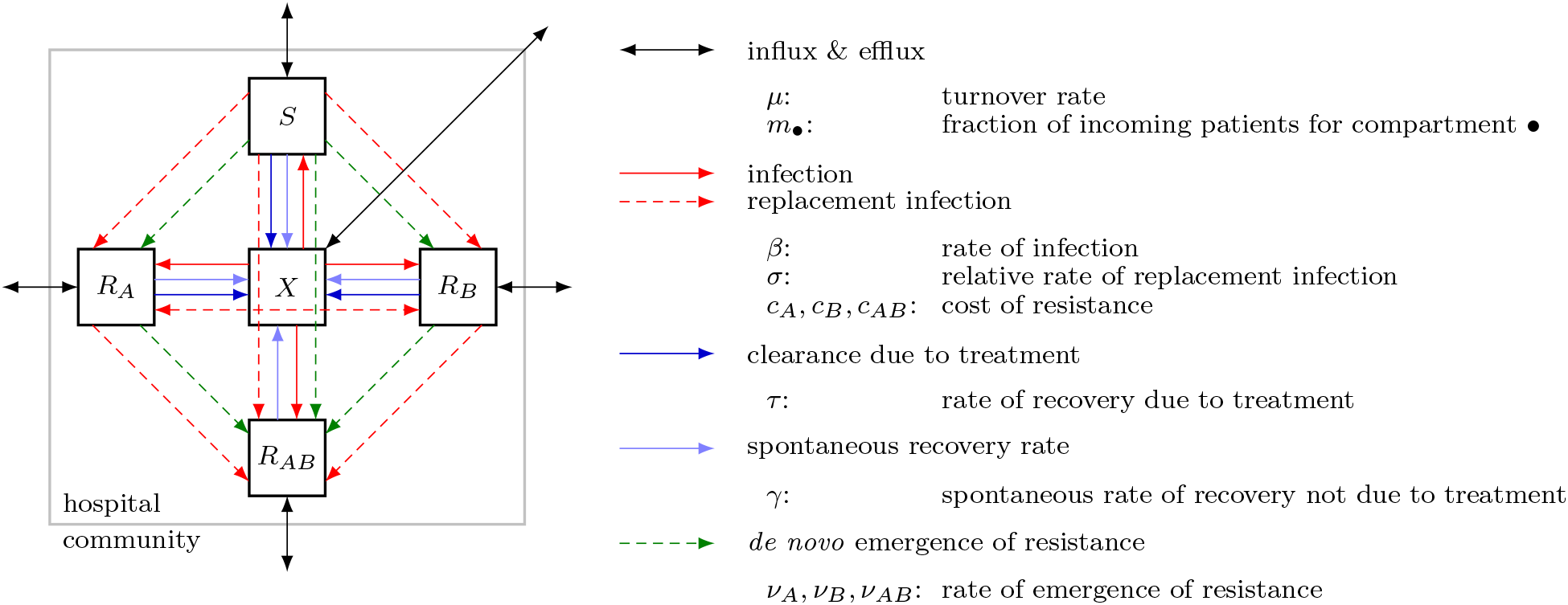
Flow diagram of the model defined by Eq. (1). The diagram shows the five model compartments and the processes that lead to patient flow between them. Solid lines describe epidemiological processes; processes marked by dashed lines involve evolution and competition between strains at the within-host level. Diagram adapted from [2] and reused with minor modification from [5].

New patients get admitted to the hospital at a total rate of *μn*_tot_. They can be uninfected or infected with either strain as given by the probability *m_•_*. Irrespective of infection status, patients leave the unit at a per-capita rate *μ*, i.e. the bacterial infection neither increases mortality nor does it require stationary treatment. *μ* is thus the turnover rate. Patients can get newly infected within the hospital. The transmission rate for the sensitive strain is *β*. The cost of resistance manifests itself in a lower transmission rate (reduction by a factor (1 − *c_•_*)). While we exclude co-infection from the model and assume that every patient is only colonized by a single strain at any time, we allow for the instantaneous replacement of infecting strains by better-adapted strains, which we term “replacement infection”. Colonization of an infected patient happens at a lower probability than colonization of an uninfected patient (reduction by a factor *σ*). The immune system clears infections at rate *γ*. A drug to which the infecting strain is susceptible leads to recovery at rate *τ*.

Finally, resistance can evolve under drug pressure. During treatment with drug A or B, resistance to the respective drug evolves at rate *ν_A_* and *ν_B_* respectively. Sensitive strains become resistant to both drugs simultaneously at rate *ν_AB_*. These rates combine mutation and fixation of the resistant strain; they also contain selection of pre-existing mutants. We assume that reversal of drug resistance within a patient due to back mutation or replacement infection through the sensitive strain is negligible.

We assume that only infected patients receive antibiotics (no prophylactic treatment). *χ_A_*, *χ_B_*, and *χ_AB_* are the fractions of infected patients that get treated with drug A, drug B, or both drugs, respectively. For combination therapy, we have *χ_AB_* = 1; for mixing, *χ_A_* = *χ_B_* = 1*/*2; for cycling, *χ_A_* = 1, *χ_B_* = 0 in periods during which drug *A* is used and *χ_A_* = 0, *χ_B_* = 1 in periods during which drug *B* is used. We always start the cycling protocol with drug A. Note that these fractions are identical for all compartments and constant in time (except for drug cycling).

Overall, we obtain the following set of ordinary differential equations (ODEs) that describe the flow between the different compartments:

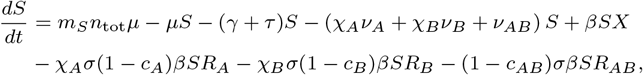

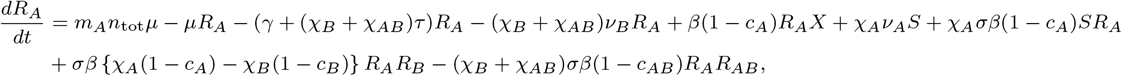

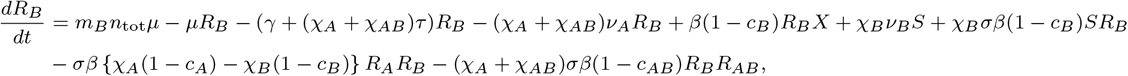

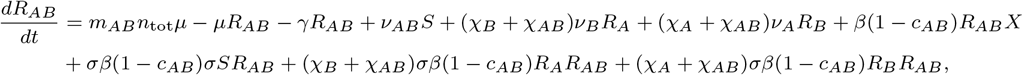

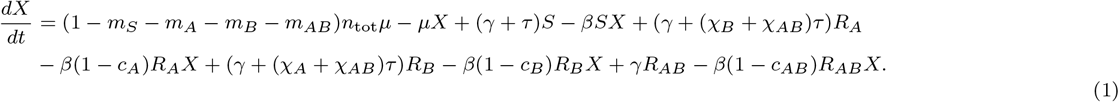

We numerically integrate Eq. (1) using Mathematica version 10.4.1.0 (Wolfram Research).

## Factors affecting the ranking of strategies

### Different optimality criteria may suggest different strategies

Some optimality criteria focus on overall treatment success (number of uninfecteds/infecteds in some time interval or at equilibrium, number of inappropriately treated patients) and others on the dynamics of resistant strains (number of patients infected with a resistant strain, emergence or spread of double resistance). It is not surprising that different treatment strategies might optimize different quantities and that a strategy might be well-suited to achieve one goal but performs poorly to achieve another [1].

In Figure 2, we compare the performance of the three strategies under three different optimality criteria (parameters and performance scores for all examples are given in SI section S2). Criterion 1 is the total number of uninfected patients within the first year 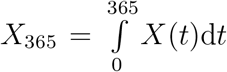 ranking the strategies as \combination therapy > mixing > cycling".Criterion 2 is the time *t_c_* by which the frequency of patients infected by the double resistant strain has reached 10% of the total number of patients. With this, we get the reverse order. Criterion 3 combines both components – number of uninfecteds and time to double resistance – in that it counts the number of uninfecteds until time 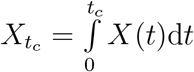 leads to the same ranking as criterion 2. Unlike criterion 1, the time period of observation differs substantially among strategies, strongly affecting the performance score.

**Figure 2:**
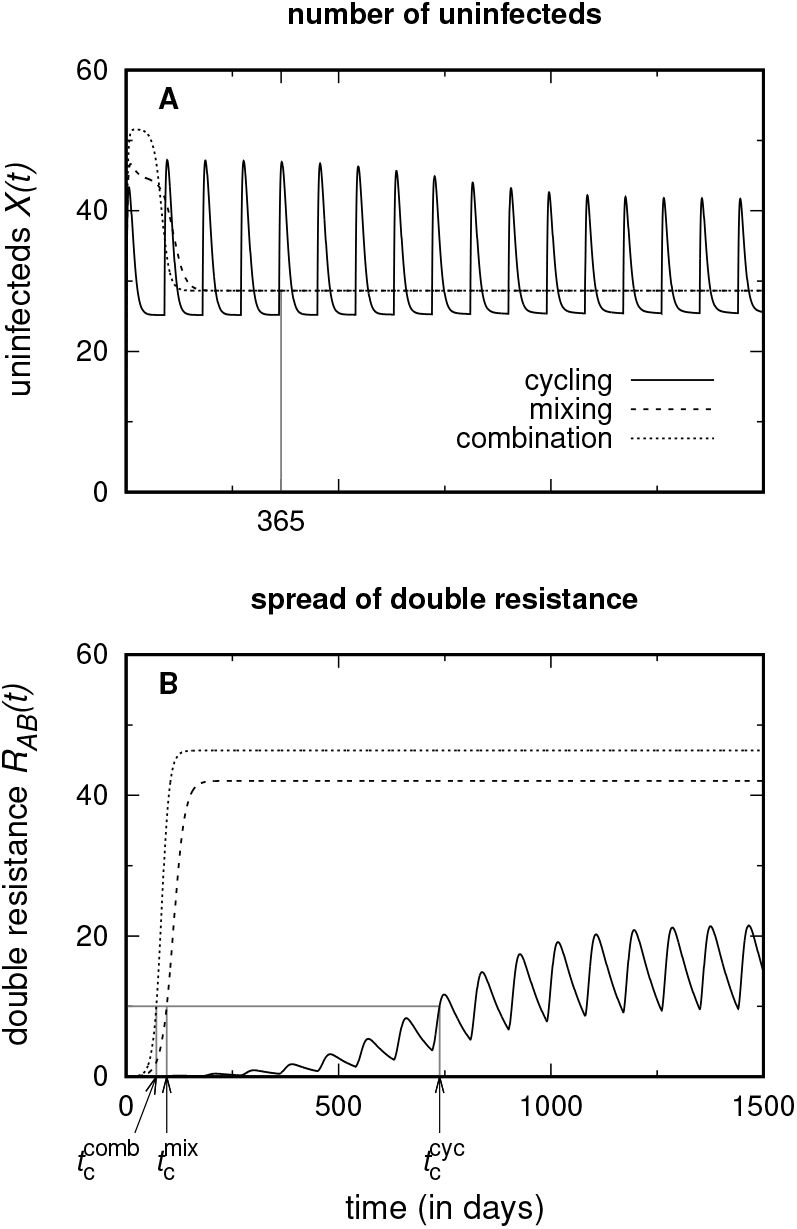
Assessment of the three treatment protocols using different optimality criteria. While the criterion “*total number of uninfected patients within the first year* “ (Panel A) ranks the strategies as “combination therapy *>* mixing *>* cycling”, the criteria “*time until the number of patients infected with the double resistant strain has reached 10% of all patients*” and “*total number of uninfected patients until this time*” (Panel B) lead to the ranking “cycling *>* mixing *>* combination therapy”. Note also that with cycling, the double resistant strain never reaches a very high frequency.

We discuss the advantages and disadvantages of the various criteria below in the discussion section.

### The model implementation may change the ordering of strategies

Following Bergstrom et al. [4], some studies use a model variant without the *R_AB_*-compartment (Figure 3A). This describes the dynamics prior to the (stochastic) emergence and establishment of double resistance. Naturally, the total number of uninfecteds in a given time interval can yield a different ranking, depending on whether an *R_AB_* compartment is included or not (Figure 4). In order to assess the risk of multiple resistance, studies based on this model variant consider the rate at which double resistance is generated from single resistant strains or the sensitive strain. While such an approach captures the time until the double resistant strain first appears, it does not make any statements about how fast it will spread through the community. Figure S3.1A considers the rate of emergence only, given by *χ_A_*(*t*)*ν_A_R_B_*(*t*) + *χ_B_*(*t*)*ν_B_R_A_*(*t*) + *ν_AB_S*(*t*) for cycling and mixing and by *ν_A_R_B_*(*t*) + *ν_B_R_A_*(*t*) + *ν_AB_S*(*t*) for combination therapy; the *R_AB_* compartment is not included into the model equations. In contrast, Figure S3.1B compares the treatment protocols with respect to the spread of double resistance, explicitly modeling the number of patients infected with the double resistant strain. The double resistant strain is generated with a higher rate under cycling than under combination therapy (and with the highest rate for mixing). However, it spreads slowest under cycling and fastest under combination therapy. This occurs because competition with the single resistant strains hampers its frequency increase under cycling. (This is at the same time another example of different optimality criteria leading to different conclusions.)

**Figure 3:**
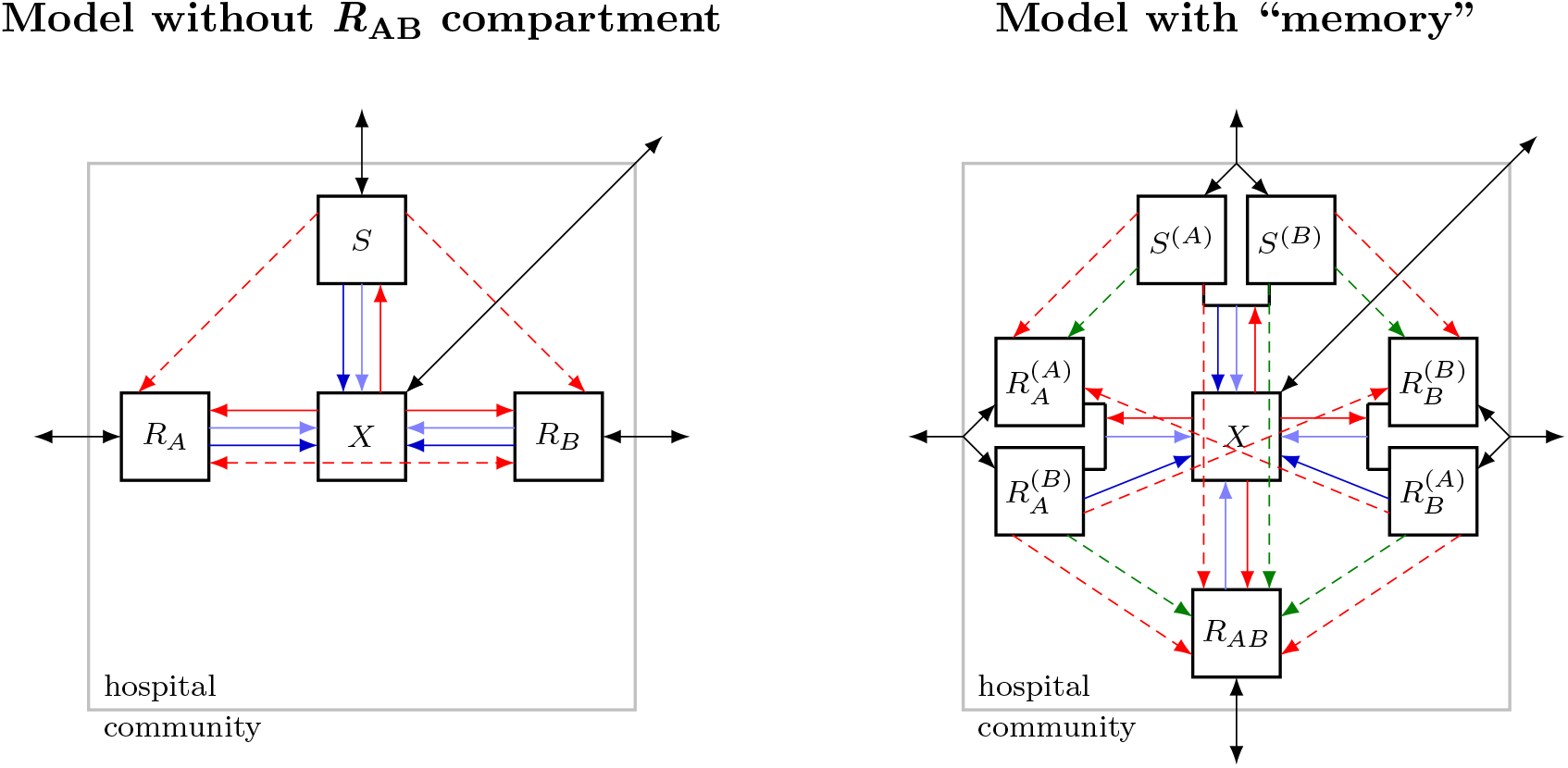
Flow diagrams of alternative model implementations. The left diagram belongs to a model without an *R_AB_* compartment (no pre-existence, no influx, and no *de novo* emergence of the double resistant strain). The right diagram reflects a model where patient groups are classified not only according to the infecting strain but also according to the drug that they receive. This model, as drawn, assumes that individual patients receive the same drug throughout their entire course of treatment. Diagrams reused with minor modification from Uecker and Bonhoeffer [5].

**Figure 4:**
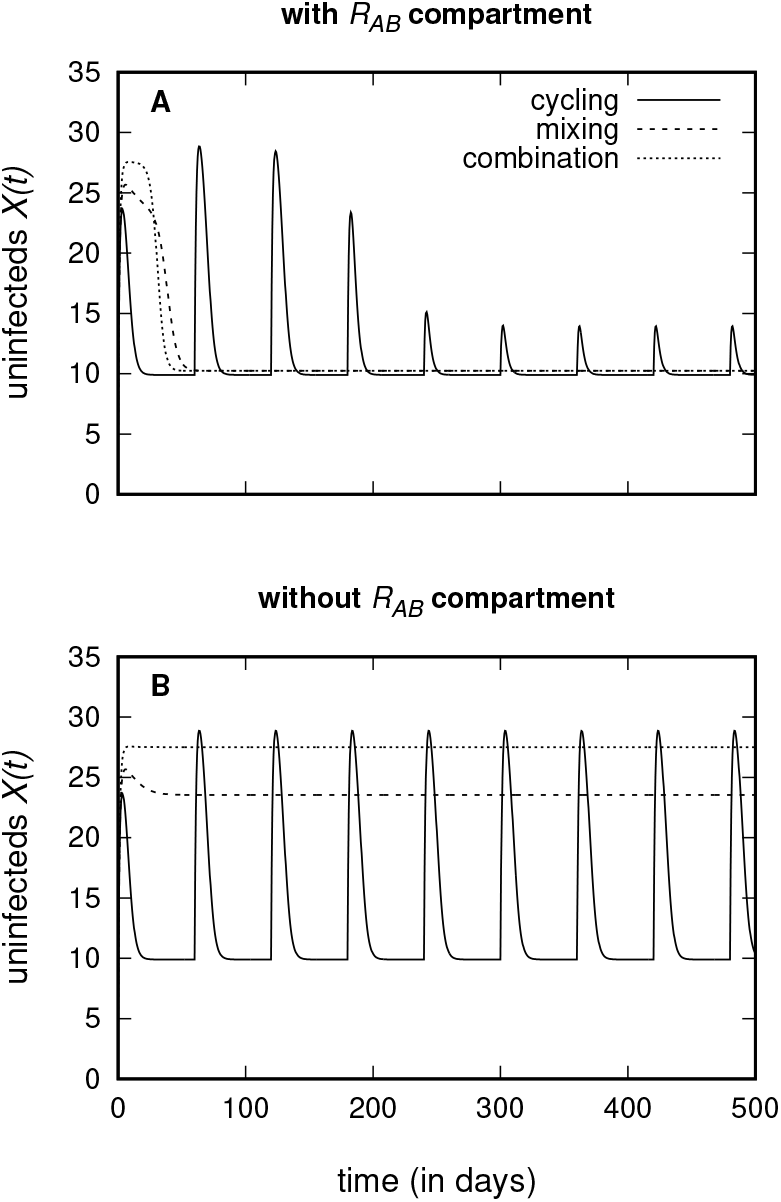
Number of uninfected patients when the model contains the *R_AB_* compartment (Panel A) and when it does not contain the *R_AB_* compartment (Panel B). For Panel A, using the number of uninfecteds within the first year as a criterion, the strategies rank “mixing ≳ cycling *>* combination therapy”. For Panel B, we obtain “combination therapy *>* mixing *>* cycling”.

Second, the model can be implemented stochastically rather than deterministically. Kouyos et al. [11] consider strategies where drugs do not get switched with a fixed period but in response to the frequency of resistance in the hospital. They find that this brings an advantage over mixing only when stochasticity is taken into account, while the difference disappears in a deterministic system.

Last, the traditional modeling approach simplifies the mixing and cycling strategies [5]. In clinical practice, patients receive the same drug throughout the entire course of treatment (unless the drug is not working but we here disregard such a drug switch), and the fractions of patients in any compartment receiving either drug are dynamic due to differential recovery rates of correctly and incorrectly treated patients. In contrast, the traditional models assume that in any compartment at any time, the fractions receiving either drug are determined by the protocol without any “memory” of which drug got assigned to individual patients. This memory can be modeled by splitting the compartments *S*, *R_A_*, and *R_B_* into two compartments each, where the new compartments not only distinguish by which strain a patient is infected but also which drug it receives (for a flow diagram, see Figure 3B). Figure S3.2 shows that this can lead to a different ranking of treatment strategies than the standard modeling approach.

### The choice of parameters and the situation prior to treatment may affect the ranking of strategies

The model as used in this study has 15 parameters (not counting the total population size *n*_tot_ that only scales the results, provided *βn*_tot_ is kept constant). More complex models accounting for more features use even more parameters. Trivially, all conclusions from numerical simulations are only valid for the chosen parameter sets (see e.g. Figure S3.3).

It has been less appreciated that not only the chosen parameter values but also the initial conditions for the ODE system (Eq. (1)) may influence the relative ranking of treatment protocols, since the number of patients in each compartment at time *t* = 0 influences the early dynamics (Figure S3.4).

## Discussion

We have highlighted a series of factors that influence the ranking of treatment strategies even though the underlying biological assumptions are (mostly) not altered. In the following, we discuss how the problems that we outlined in the previous section can be approached and suggest new modeling directions that would make the models more compatible with clinical studies.

### The optimality criterion

The choice of the optimality criterion is particularly critical for the meaningful assessment of a treatment protocol and deserves very careful consideration, both in models and clinical trials. A range of criteria are currently applied, and this can substantially influence the conclusions. Hence, which criterion should be used? If the goal is to enhance our understanding of the evolutionary dynamics, any of the above listed criteria could be insightful and meaningful. If the goal is to foster clinical trials or to come to practically relevant conclusions, answering this question is difficult. Essentially, three factors need to be taken into account: the clinical benefits (e.g. clearance of the infection) and costs (e.g. side effects) and economic costs. The modeling framework directly only targets the first of these factors. Moreover, a good criterion that aims at applicability should not only be theoretically convincing but moreover needs to be suitable to serve as a clinical endpoint.

Generally, we advocate to use a criterion that aims at maximising overall treatment success rather than to focus on resistance evolution. This is because eventually, we are not interested in the evolution of resistance per se – resistance could be avoided by simply not treating anybody – but rather in its harmful consequence, which is to prevent (rapid) patient recovery.

From the criteria listed in Table 1, hospital-wide disease prevalence assesses overall treatment success, and its changes upon installment of a new antibiotic treatment protocol are measurable. Two criteria that have not received much attention in the mathematical literature so far are the mortality rate and the length of hospital stay, which are however criteria that are used in order to measure the health burden of resistant infections [but see 1]. Following Bergstrom et al. [4], many models assume that the efflux rate is independent of the infectious status, and both criteria are hence inherently meaningless in these models. Some models, in the tradition of Bonhoeffer et al. [3], allow for differential efflux rates of infecteds and uninfecteds. However, they do not distinguish between discharge and death (partially because they do not consider the dynamics of a hospital but of a general community, where discharge of recovereds does not occur). While it might not always be possible to clearly assign a fatal outcome to a nosocomial disease (rather than to the original cause of hospitalisation), changes in the total mortality rate after a treatment protocol became implemented can be easily measured. Likewise, the length of hospital stay can be readily measured. It is important to note that neither of these three criteria– disease prevalence, mortality rate, length of hospitalisation – is sufficient on its own but an assessment of the mortality rate should be combined with an assessment of disease prevalence or the duration of hospitalisation in order to take both outcomes of inappropriate treatment (death and prolonged illness) into account. A caveat with all three criteria is that one needs to choose a time period during which their performance is assessed. At short time scales, the outcome is dominated by the transient behavior following installment of the new protocol. In contrast, with a long observation period, the equilibrium dynamics determine the ranking. Depending on the chosen time frame, conclusions might hence differ, and optimally, both the short-term and the long-term performance should be assessed.

While we think that treatment success (in whatever way assessed) is a better measure of quality than reduced resistance, monitoring resistance under the various treatment strategies is by no means irrelevant for their evaluation, neither in modeling nor in clinical studies. One reason for this is that antibiotics may be abandoned when resistance reaches a threshold. I.e. even if a treatment strategy reduces overall treatment failure compared to another protocol, it might lead to the earlier removal of a certain antibiotic. Given the decelerated discovery of new antibiotics, this will reduce treatment options. Moreover, the alternative antibiotic might have more severe side effects or might be substantially more expensive. A priori, we think that criteria that incorporate the spread of resistance are more relevant than those purely quantifying the rate of emergence. Moreover, the strong stochasticity makes the first appearance of resistance an unsuitable criterion for clinical trials. For clinical trials, a criterion that integrates information from a time period is more robust than a criterion that relies on a single incidence or time point.

A criterion that explicitly combines the efficacy of treatment and the spread of resistance is provided by the number of uninfected patients until the frequency of (double) resistance has reached a threshold. On the one hand, this is an attractive measure, in particular for situations in which the antibiotic is abandoned when a resistance threshold is reached. On the other hand, it is uninformative in that it entirely ignores what happens after that point of time, and this time is potentially very different for different strategies. In particular, this time is infinite in the absence of treatment, implying that treating no-one achieves a perfect score, given that the number of uninfecteds at equilibrium is non-zero. Moreover, such a composite criterion makes it more difficult to understand the underlying dynamics. For clinical trials, it is again problematic since the prevalence of resistance is most likely subject to strong fluctuations, at least if measured in small units such as in a single hospital ward. Hence, the point of time at which resistance crosses the threshold is subject to stochastic variation, adding considerable extra noise and uncertainty to the data.

Since the multifacetedness of the problem does not allow us to pin down one “correct” universally applicable criterion, how can we still make progress? Applying more than one criterion (ideally targeting treatment success and resistance) seems to be a sensible approach and has also been done in several studies in the past (see Table 1). Even if no strategy is the best under all aspects, it is helpful to know under which criterion it is the best. This knowledge makes it possible to make an informed decision in specific cases, depending on which criterion seems to be the most important one under the given circumstances. E.g., for infections, where the mortality rate is high unless appropriate treatment is initiated immediately, good performance under a criterion that evaluates overall treatment success is more relevant than good performance under a criterion that focuses on the emergence of resistance. This is particularly true if a large number of antibiotics are available for this particular bacterial species. In contrast, for an infection that takes a mild course in most patients even if treated late, it might be preferable to control resistance as well as possible in order to maintain efficient treatment options with mild side effects for rare severe cases. As a side remark, to put optimality scores into context, it would be interesting to compare potential benefits of multi-drug strategies to improvements achieved by other means, such as a reduction in transmission.

### The model implementation

It is clear that excluding the possibility of double resistance from the analysis neglects a core aspect of the problem. Yet, whether the model best contains an *R_AB_* compartment or not, depends on the question to be answered. The *de novo* emergence of double resistance is a highly stochastic process. If the double resistant strain is initially absent, there is a phase before it is generated (or is brought into the hospital from the outside) and starts spreading. Models without the *R_AB_* compartment allow to assess the performance of protocols during this phase and to estimate its length. Deterministic models incorporating an *R_AB_* compartment ignore this phase but allow to study the spread of double resistance.

Generally, a deterministic model implementation assumes that all patient groups are large enough to neglect stochastic fluctuations. However, many models focus on treatment strategies in hospital wards, which normally only accommodate a relatively small number of patients. Even a hundred patients is not a large population size if “large” refers to the negligibility of stochasticity. Stochastic models are therefore a priori more appropriate than deterministic implementations. Stochasticity is very high in clinical trials, making it hard to arrive at robust conclusions. Here, stochastic models can help to assess which conclusions can and cannot be drawn in the face of randomness and give a sense of the scale at which clinical studies would need to be performed. However, the role and importance of certain parameters and processes can presumably be well assessed using deterministic models.

Last, the traditional modeling approach classifies compartments only according to the infecting strain and does not take the administered drug into account as well [5]. This approach disregards the associations that build up between the infecting strain and the drug used. (These associations build up since patients treated with the wrong drug recover more slowly.) This simplified model is often a good approximation, leading to similar predictions. However, awareness of the simplification seems important.

### The model analysis

The models are challenging to analyse for two reasons. First, due to the model complexity, most studies rely on numerical simulations rather than on analytical approximations. It is hence not possible to read off general results from an analytical solution. Second, the parameter space is very large and parameter estimates are lacking.

Parameter sensitivity and structural sensitivity tests can alleviate the problem (see Tepekule et al. [2] for a recent study that implements random sampling of parameters, linear discriminant analysis, and particle swarm optimization to systematically explore the parameter space; for an insightful discussion of structural sensitivity in between-host models of antibiotic resistance, see Spicknall et al. [20]). Importantly, this includes to allow for asymmetry between the two single resistant strains and for unequal use of the two drugs, e.g. a drug ratio other than 50:50 for the mixing strategy [7]. It is also important to investigate by which margin strategies differ from each other (and from mono-drug therapies, which sometimes even outperform multi-drug strategies, [2]). If differences are small, a ranking may be meaningless.

For the ranking of strategies, a numerical analysis is required to be able to allow for sufficient model complexity. For a more fundamental understanding of the model behavior, e.g. for understanding the reasons why a strategy performs better or worse than expected, an analytical treatment of simplified models or limiting cases may provide further insight.

### Important model extensions

The modeling framework allows, of course, for innumerous extensions but we will only discuss three potential directions here. The first set of suggestions is motivated by the above discussion of the optimality criterion. As already pointed out, current models typically either assume that patients leave the hospital or die independent of their infectious status or allow for differential efflux rates but do not differentiate the cause of efflux (i.e., discharge or death) and make no use of this feature. However, empirical therapy is especially important for critically ill and immuno-compromised patients where the bacterial infection constitutes a true burden on their health and a risk to their life. It is hence very likely that the choice of treatment – effective or ineffective – influences (1) the duration of hospitalisation and (2) the survival chances of the patient. It therefore seems highly relevant to focus on rates of discharge and death that depend on the infectious status of the patient. In this setting, the length of stay in the hospital and the mortality rate could be used as a measure of success for a treatment strategy. To assess these quantities in a deterministic framework, it is straightforward to complement the current ODE system by two further compartments, one for successfully treated and discharged (former) patients and one for the deceased. The size of the former is closely linked to the length of hospitalisation; the size of the latter is closely linked to the mortality rate.

Second, nested models that describe both the within-host dynamics of the bacteria and the epidemiological dynamics could help to gain a more realistic picture. In the current epidemiological models, the transition rates (recovery rate, emergence of resistance etc.) are modeled by independent parameters. However, these rates are in reality coupled and explicit modeling of the within-host dynamics would account for these inter-dependencies. Of course, the approach would require us to make assumptions about the parameters at the within-host level that are equally unknown as the parameters at the between-host level, hence leading to considerable uncertainty about the appropriate model and its parametrization. Nested models should not replace but complement the current simpler models.

In a recent study, Beardmore et al. [1] took a step forward with respect to both of the points discussed here. In a briefly presented model, the bacterial load within individual patients is tracked, and discharge of patients depends on their bacterial load, making the length of hospital stay variable (there is no mortality in the model). The duration of hospitalisation is used to assess the performance of the two treatment strategies cycling and mixing. Further development of models along these lines could greatly enhance our understanding of the prospects of antibiotic treatment protocols.

Last, nosocomial infections are often caused by commensal bacteria. They may stem either from the patient’s own flora and may have already been present at admission or acquired asymptomatically within the hospital. Or they may be acquired as pathogens from other patients. These different routes of infection and ways of transmission are not accounted for by most modeling studies. Most studies follow the approach of traditional epidemic models for the spread of obligate pathogens. E.g., transition from the *X* to any infected compartment is only possible through infection from infected patients rather than through self-infection from own commensals. Rethinking the models on the background of commensal bacteria as agents of infection seems to be important in order to build a framework that is fully consistent with its objectives. In this context especially, the consequences of horizontal gene transfer should be considered. Clinically relevant resistance is often encoded on plasmids. Yet, how this influences the effectiveness of cycling, mixing, or combination therapy has only been touched upon [4].

### Concluding remarks

Even if we might have preferred to identify one strategy as the universally best, the complex picture that has emerged from mathematical models so far should be appreciated as a result in its own right. It should also be appreciated that it is the result of a scientific discourse and a development brought forward by a series of articles. It was by no means clear a priori that no simple answer exists.

The focus of this article is on the role of mathematical models in the assessment of antibiotic treatment protocols and on ways to improve their contribution. However, a joint effort is necessary to arrive at good solutions. The theoretical work reviewed in this article is not discussed here in the context of clinical studies, while such a connection could lead to more potent studies. This would be particularly valuable if through sequencing and the analysis of sequence data, the pathway of resistance could be traced back along patients in order to distinguish *de novo* acquired from transmitted resistance. Besides models and clinical studies, a third tool, which has surprisingly been ignored so far, is *in vitro* experimental evolution. While evolution experiments are widely used to investigate the evolution and maintenance of antibiotic resistance and are also used to study the effect of combining antibiotics, population-wide treatment strategies have to our knowledge not been simulated in the lab. As an intermediate between models and clinical trials, they could help to close the gap between theoretical and clinical studies in the future.

## Acknowledgements

We thank Stephanie Fingerhuth and Andrew Read for helpful discussions and Sally Otto for valuable comments on the manuscript.

